# The genetic architecture of local adaptation I: The genomic landscape of foxtail pine (*Pinus balfouriana* Grev. & Balf.) as revealed from a high-density linkage map

**DOI:** 10.1101/011106

**Authors:** Christopher J. Friedline, Brandon M. Lind, Erin M. Hobson, Douglas E. Harwood, Annette Delfino Mix, Patricia E. Maloney, Andrew J. Eckert

## Abstract

Explaining the origin and evolutionary dynamics of the genetic architecture of adaptation is a major research goal of evolutionary genetics. Despite controversy surrounding success of the attempts to accomplish this goal, a full understanding of adaptive genetic variation necessitates knowledge about the genomic location and patterns of dispersion for the genetic components affecting fitness-related phenotypic traits. Even with advances in next generation sequencing technologies, the production of full genome sequences for non-model species is often cost prohibitive, especially for tree species such as pines where genome size often exceeds 20 to 30 Gbp. We address this need by constructing a dense linkage map for foxtail pine (*Pinus balfouriana* Grev. & Balf.), with the ultimate goal of uncovering and explaining the origin and evolutionary dynamics of adaptive genetic variation in natural populations of this forest tree species. We utilized megagametophyte arrays (*n* = 76–95 megagametophytes/tree) from four maternal trees in combination with double-digestion restriction site associated DNA sequencing (ddRADseq) to produce a consensus linkage map covering 98.58% of the foxtail pine genome, which was estimated to be 1276 cM in length (95% CI: 1174 cM to 1378 cM). A novel bioinformatic approach using iterative rounds of marker ordering and imputation was employed to produce single-tree linkage maps (507–17 066 contigs/map; lengths: 1037.40– 1572.80 cM). These linkage maps were collinear across maternal trees, with highly correlated marker orderings (Spearman’s *ρ >* 0.95). A consensus linkage map derived from these single-tree linkage maps contained 12 linkage groups along which 20 655 contigs were non-randomly distributed across 901 unique positions (*n* = 23 contigs/position), with an average spacing of 1.34 cM between adjacent positions. Of the 20 655 contigs positioned on the consensus linkage map, 5627 had enough sequence similarity to contigs contained within the most recent build of the loblolly pine (*P. taeda* L.) genome to identify them as putative homologs containing both genic and non-genic loci. Importantly, all 901 unique positions on the consensus linkage map had at least one contig with putative homology to loblolly pine. When combined with the other biological signals that predominate in our data (e.g., correlations of recombination fractions across single trees), we show that dense linkage maps for non-model forest tree species can be efficiently constructed using next generation sequencing technologies. We subsequently discuss the usefulness of these maps as community-wide resources and as tools with which to test hypotheses about the genetic architecture of adaptation.

## Introduction

Evidence for adaptive evolution among populations of plants is commonly documented at the phenotypic and molecular levels (Kawecki and Ebert 2004; Pannell and Fields 2013), as such some of the best examples of adaptive evolution within lineages come from the field of plant genetics (e.g., Antonovics and Bradshaw 1970). Despite this evidence, relatively little work has focused explicitly on the genomic organization of loci contributing to these patterns (Hoffmann and Riesberg 2008), which likely stems from a lack of genomic resources for plants relative to animals. Adaptive evolution has been extensively documented for forest trees, especially conifers, with many instances of local adaptation clearly documented over the past century (White et al. 2007; Neale and Kremer 2011). Despite great advances in experimental technology, empirical focus has remained almost fully on the number, effect size, type, and interactions among loci contributing to adaptive evolution (Neale and Kremer 2011; Alberto et al. 2013). A thorough examination of the genetic architecture of fitness-related traits, however, should also include an examination of the genomic organization of the loci contributing to trait variation. We leverage this idea in the first of a series of papers dissecting the genetic architecture of fitness-related traits in a non-model conifer species, foxtail pine (*Pinus balfouriana* Grev. & Balf.).

The genomic organization of loci contributing to variation in fitness-related traits would follow naturally from the production of a sequenced genome (i.e., a physical map). For many taxa, especially those with small to modest genome sizes, this is monetarily and computationally feasible using next-generation DNA sequencing technologies (Koboldt et al. 2013). For taxa with large or complex genomes, however, even the advent of next generation DNA sequencing does not solve the complexity and cost hurdles associated with the production of a finished genome sequence. Conifers have large and complex genomes (Murray 1998; Ahuja and Neale 2005), with estimated average genome sizes in *Pinus* in the range of 20 Gbp to 30 Gbp. Several genome projects, each of which involves large consortia, are underway or have been completed (Mackay et al. 2012). Even these efforts often initially result in limited information, however, as for example the current assemblies of the Norway spruce (*Picea abies* L.) and loblolly pine (*Pinus taeda* L.) genomes contain millions of unordered contigs with average sizes in the thousands of base pairs (Nystedt et al. 2013; Neale et al. 2014). An alternative, but not mutually-exclusive, approach to describing the genome of an organism is that of linkage mapping. In this approach, genetic markers are ordered through observations of recombination events within pedigrees. This approach dates to the beginning of genetics and the logic has remained unchanged since the first linkage maps were created in *Drosophila* (Sturtevant 1913).

Renewed interest in linkage maps has occurred for two reasons. First, linkage maps are often used to order contigs created during genome sequencing projects (Mackay et al. 2012; Martinez-Garcia et al. 2013). In this fashion, linkage maps are used to help create larger contigs from those generated during the assembly. It is these larger contigs that create the utility that most practicing scientists attribute to genome sequences. Second, linkage maps are easy to produce and provide a rich context with which to interpret population and quantitative genetic patterns of variation (e.g., Eckert et al. 2010b, a, 2013; Yeaman 2013). They can also be used to test explicit hypotheses about the organization of loci contributing to adaptive evolution. For example, Yeaman and Whitlock (2011) developed theoretical predictions about the genomic organization of loci underlying patterns of local adaptation as a function of gene flow, so that loci contributing to local adaptation have differing spatial structure within genomes as a result of differing regimes of gene flow. The relevant scale (*sensu* Houle et al. 2011) in these mathematical formulations is that of recombinational distance among loci, so that when matched with an appropriate study system, linkage maps provide the impetus to test basic evolutionary hypotheses. In this context, future additions of finished genome sequences would add to the interpretation of results.

Construction of linkage maps have a long history within forest genetics, mostly through their use in quantitative trait locus mapping (Ritland et al. 2011). Conifers in particular are highly amenable to linkage mapping, with approximately 25 different species currently having some form of linkage map completed (see Table 5-1 in Ritland et al. 2011). Much of the amenability of conifers to linkage mapping stems from the early establishment of breeding populations in economically important species and from the presence of a multicellular female gametophyte (i.e., the megagametophyte) from which the haploid product of maternal meiosis can be observed (Cairney and Pullman 2007). Indeed, many of the first linkage maps in conifers were generated from collections of megagametophytes made from single trees (Tulsieram et al. 1992; Nelson et al. 1993; Kubisiak et al. 1996). Continued advancements in genetic marker technologies have facilitated rapid development of linkage maps across a diversity of species (e.g. Achere et al. 2004; Kang et al. 2010; Martinez-Garcia et al. 2013). The development of biologically informative markers for non-economically important conifers, however, is hampered by production costs associated with the creation of characterized genetic markers (i.e., those with a known DNA sequence and/or function). The majority of this cost is in the two-step approach needed to generate biologically meaningful markers: polymorphism discovery via DNA sequencing followed by genotyping of those polymorphisms (cf., Eckert et al. 2013). As a result, the vast majority of linkage maps outside of economically important species are created with uncharacterized genetic markers (e.g., Travis et al. 1998). Much of the knowledge about the genetic architecture of fitness-related traits, outside of a handful of well studied conifer species, therefore, encompasses the number and effect size of uncharacterized genetic markers (Ritland et al. 2011). Cost restrictions, however, have largely disappeared. It is now feasible to jointly discover polymorphisms and genotype samples using high-throughput DNA sequencing approaches, such as restriction site associated DNA sequencing (RADseq; e.g., Peterson et al. 2012).

The generation of linkage maps from RADseq data is a complex endeavor due to the inherent stochasticity and error prone nature of these data. Recent examples in several crop species highlight the difficulties that must be overcome with respect to missing data and errors in calling polymorphic sites and the resulting genotypes (Pfender et al. 2011; Ward et al. 2013). Despite these difficulties, RADseq has been successively applied to samples taken from natural populations of non-model conifer species (Parchman et al. 2012), but has yet to be applied to linkage mapping in these species. An exploration of these methods to linkage mapping in the large and complex genomes of conifers is thus warranted. Here, we take this approach using megagametophyte arrays from four maternal trees of foxtail pine to generate maternal linkage maps. There are currently no published linkage maps for this species, which is only distantly related to loblolly pine (Eckert and Hall 2006), nor any within the subsection *Balfourianae*. We subsequently discuss the utility of our inferred linkage maps to tests of evolutionary theory addressing local adaptation and its genetic architecture.

## Materials and Methods

### Focal species

Foxtail pine is a five needle species of *Pinus* classified into subsection *Balfourianae*, section *Parrya*, and subgenus *Strobus* (Gernandt et al. 2005). It is one of three species within subsection *Balfourianae* (Bailey 1970) and generally is regarded as the sister species to Great Basin bristlecone pine (*P. longaeva* D. K. Bailey; see Eckert and Hall 2006). The natural range of foxtail pine encompasses two regional populations located within California that are separated by approximately 500 km: the Klamath Mountains of northern California and the Sierra Nevada of southern California (Figure 1). These regional populations diverged approximately one million years ago (mya), with current levels of gene flow between regional populations being approximately zero (Eckert et al. 2008). Within each regional population, levels of genetic diversity and the degree of differentiation among local stands differ, with genetic diversity being highest in the southern Sierra Nevada stands and genetic differentiation being the highest among the Klamath stands (Oline et al. 2000; Eckert et al. 2008). These two regional populations have also been recognized as distinct subspecies based on numerous quantitative traits, with *P. balfouriana* subsp. *balfouriana* located in the Klamath region and *P. balfouriana* subsp. *austrina* located in the southern Sierra Nevada mountains (Mastrogiuseppe and Mastrogiuseppe 1980). The two regional populations of foxtail pine thus represent a powerful natural experiment within which to examine the genomic organization of loci contributing to local adaptation. The first step in using this system to test evolutionary hypotheses is the production of a dense linkage map (cf., Pannell and Fields 2013).

**Fig. 1.**
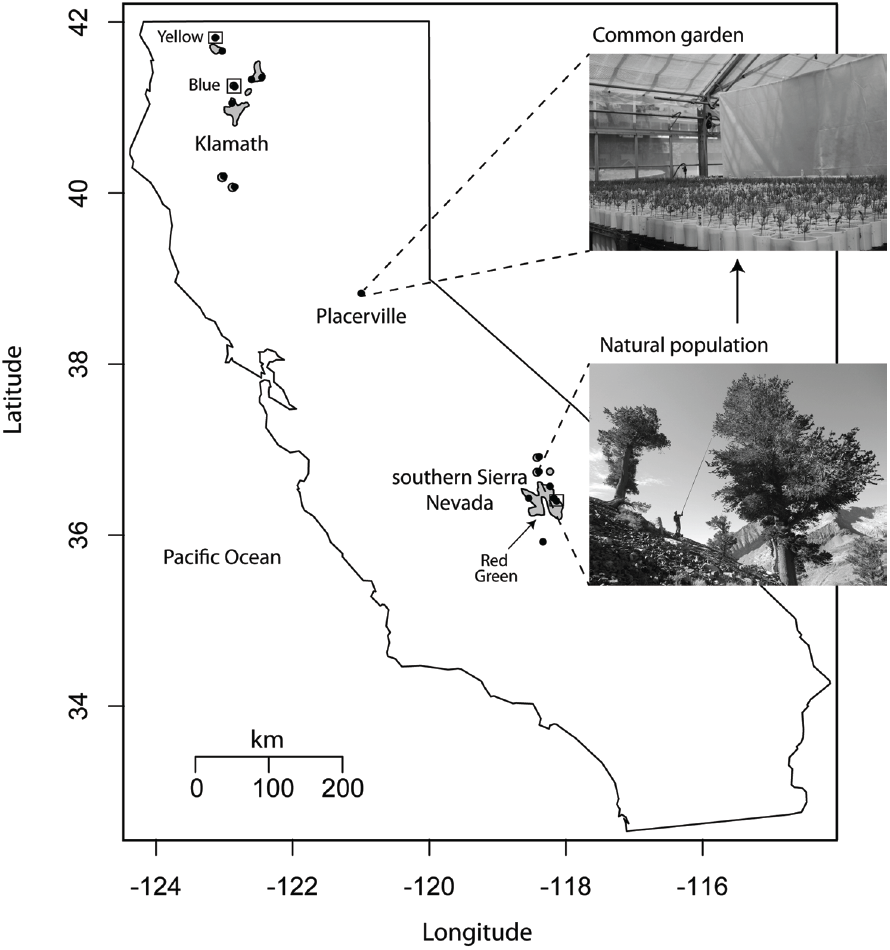
Geographical locations of foxtail pine samples used to construct a common garden located in Placerville, CA. Circles denote the 15 unique locations from which 4 to 17 maternal trees were sampled. Circles enclosed in squares denote locations from which maternal trees used in linkage mapping were sampled. Photo credits: lower: T. Burt; upper: A. Delfino Mix

### Sampling

Seed collections from 141 maternal trees distributed throughout the natural range of foxtail pine were obtained during 2011 and 2012. Of these 141 maternal trees, 72 were sampled from the Klamath region, while 69 were sampled from the southern Sierra Nevada region. These 141 families were divided among 15 local stands (*n* = 4 trees/stand to 17 trees/stand), with eight stands in the Klamath region and seven stands in the southern Sierra Nevada. Approximately 50 seeds were germinated from each seed collection and 35 of those 50 seedlings were planted in a common garden located at the USDA Institute of Forest Genetics, Placerville, California (Figure 1). The common garden was established using a randomized block design and involved three separate plantings of seeds spanning approximately one year (June 6, 2012 until May 20, 2013). Four of the 141 maternal trees were selected at random (*n* = 2 from the Klamath region and *n* = 2 from the southern Sierra Nevada) for linkage analysis. Libraries were color-coded and are referred to as red (southern Sierra Nevada), green (southern Sierra Nevada), blue (Klamath), and yellow (Klamath). For each of these trees, 75 to 100 seeds were germinated and planted in the common garden. Upon germination, haploid megagametophyte tissue was rescued from each seedling, cleaned by removing soil and other extraneous materials with water, and stored for further analysis in 1.5 mL Eppendorf tubes at *−*20 ◦C.

### Library Preparation and Sequencing

Total genomic DNA was isolated from each rescued megagametophyte using the DNeasy 96 Plant kit following the manufacturer’s protocol (Qiagen, Germantown, MD). RADseq (Davey and Blaxter 2010; Parchman et al. 2012; Peterson et al. 2012) was used to generate a genome-wide set of single nucleotide polymorphism (SNP) markers for linkage mapping following the protocol outlined by Parchman et al. (2012). In brief, this protocol is a double-digestion, RADseq (ddRADSeq) approach based on digestion of total genomic DNA using EcoRI and MseI followed by single-end sequencing on the Illumina HiSeq platform. Single-end sequencing was chosen for reasons related to cost. Paired-end sequencing would have improved the reference assembly, which would have likely improved construction of the linkage map. Since the insert size we selected would have resulted in non-overlapping reads from each end, the improvement to genotype calls is unclear. Following digestion, adapters containing amplification and sequencing primers, as well as barcodes for multiplexing, were ligated to the digested DNA fragments. We chose to multiplex 96 samples using the barcodes available from Parchman et al. (2012). One of these samples, per set of 96, was a pseudo-diploid constructed by pooling five megagametophytes sampled from the same maternal tree, although there is a probability of 0.5^4^ = 0.0625 that the genotype for any given SNP will be mistakenly called homozygous due to the five megagametophytes all being of the same allele (see Morris and Spieth 1978). These barcodes are a mixture of 8 bp, 9 bp, and 10 bp tags that differ by at least four bases. Following ligation, successfully ligated DNA fragments were amplified using PCR and amplified fragments were size selected using gel electrophoresis. We selected fragments in the size range of 400 bp (300 bp to 500 bp) by excising and purifying pooled DNA from 2.5% agarose gels using QIAquick Gel Extraction Kits (Qiagen). Further details, including relevant reagents and oligonucleotide sequences, can be found in File S1. All DNA sequencing was performed on the Illumina HiSeq 2000 or 2500 platform at the VCU Nucleic Acids Research Facility (http://www.narf.vcu.edu/).

### DNA Sequence Analysis

There are multiple steps involved with the processing of raw DNA sequence reads into a set of SNP genotypes that are useful for linkage mapping: (1) quality control, filtering, and demultiplexing, (2) assembly to generate a reference sequence for mapping reads, (3) mapping of reads to call SNPs and genotypes for each sample, and (4) filtering of SNPs and the resulting genotypes for data quality and biological meaning.

DNA sequence reads were demultiplexed into sample-level fastq files, following quality control and filtering. The filtering pipeline was adapted from Friedline et al. (2012). Briefly, reads containing any N beyond the first base were excluded, however, reads having N as the first base were shifted by one base to the right to exclude it (i.e, a read starting with NTGC would become a read starting with TGC). Additional quality filtering ensured that all reads in the resulting set for downstream processing had a minimum average quality score of 30 over 5-bp sliding windows and that not more than 20 % of the bases had quality scores below 30. Reads passing the quality control steps were demultiplexed into sample-specific fastq files by exact pattern matching to known barcodes. Reads that did not match a known barcode were excluded.

The individual with the largest number of reads across all four maternal trees was assembled using Velvet (Zerbino, version 1.2.10), with hash length (*k*) optimized using parameter sweeps of *k* through the contributed `VelvetOptimiser` (http://www.vicbioinformatics.com, version 2.2.5) script (for odd *k* on *k* = [19, 63]). Assembly robustness was evaluated in each case using the LAP likelihood framework (Ghodsi et al. 2013), version 1.1 (svn commit r186) following mapping of the original reads to the assembly with Bowtie2 (Langmead and Salzberg 2012) (`--local --very-sensitive-local`). The assembly with the maximum likelihood value was chosen as the reference for SNP calling.

SNPs were called for all individuals against the reference using the following methodology. First, reads were mapped to the reference with Bowtie2 (`--local --very-sensitive-local`). These resulting sam files were converted to their binary equivalent (e.g., bam) using samtools version 0.1.19 (`view, sort, index`) (Li et al. 2009). SNPs were called using bcftools and filtered using vcfutils to exclude SNPs with less than 100x coverage. The resulting variant call files (vcf) were further processed using vcftools version 0.1.11 (Danecek et al. 2011) to remove indels, exclude genotype calls below a quality threshold of 5, and output as a matrix (`--012`) the haploid genotype of each megagametophyte for each SNP.

We used several thresholds to filter called SNPs for linkage mapping. First, we excluded SNPs using a *χ*^2^ test of homogeneity against an expectation of 1:1 segregation. This segregation pattern was expected because the maternal tree had to be a heterozygote to detect a SNP, and Mendel’s first law guarantees that the segregation ratio for this SNP should be 1:1. Significance of each test was assessed using a Bonferroni-corrected significance threshold of *α* = 0.05, where *α* was corrected using the number of SNPs tested. As reads from each family were mapped against a single reference assembly, we performed the *χ*^2^ test and corrections on a family-wise basis. Second, for each family, we filtered the resulting SNPs based on the genotype of the pseudo-diploid sample in that family so as to keep only those SNPs where the pseudo-diploid was either 1) called a heterozygote or 2) had a missing genotype call. Lastly, we filtered the resulting SNPs so as keep only those that had a minimum of 5 genotype calls for each of the alternate alleles. These filtering steps were taken to minimize the presence of genotyping errors arising from technical (e.g., read mapping and alignment) and biological (e.g., paralogy) reasons. Previous research within conifer genomes has documented to the presence of a large number of paralogues (Keeling et al. 2008; Nystedt et al. 2013; Neale et al. 2014). Although we did not explicitly quantify the degree of paralogy consistent with our data, the filters used during the analysis of DNA sequence reads should flag paralogous loci preferentially. The resulting subset of SNPs was then used as the input to linkage analysis.

### Linkage Analysis

The production of a linkage map requires three main steps: (1) calculation of pairwise distances between all pairs of loci, (2) clustering (i.e., grouping) of loci based on these pairwise distances, and (3) ordering of loci within each cluster (Cheema and Dicks 2009). A variety of software packages exist to carry out these steps (e.g., Van Ooijen 2011). Traditional software packages for linkage mapping, however, are not amenable to large amounts of missing data and frequent errors in genotype calls. The former causes issues with all aspects of analysis, while the latter primarily affects the genetic distances between markers (Hackett and Broadfoot 2003; Cartwright et al. 2007). We thus followed the approach of Ward et al. (2013), which was designed specifically for RADseq data.

In brief, this method can be described as follows. Pairwise distances were estimated and loci were clustered using a custom R script (R Core Team 2013). We used MSTmap (Wu et al. 2008a) to infer marker order and Maskov (Ward et al. 2013) to impute and correct genotypes. The algorithms available in MSTmap can also be used to impute and correct genotype errors (see Wu et al. 2008a), but the amount of missing data and putative genotyping errors in our RADseq data far surpassed those used to develop this software. These two programs were used in an iterative fashion. MSTmap was used initially to order markers, which was followed by the use of Maskov to impute and correct putative genotype errors conditional on this initial marker ordering. A last round of ordering was performed using MSTmap conditional on the imputed and error corrected genotype data. This general schema was followed for each of the four maternal trees independently.

The relevant pairwise distance for linkage mapping in our haploid case is defined as the probability of observing a recombination event between two haplotypes. This probability can be calculated for a set of biallelic loci using the Hamming distance (*d*_*i, j*_). The Hamming distance is the number of differences separating two binary strings (Hamming 1950), which are in this case, the haploid genotypes for a set of two megagametophytes. This distance, scaled by the number of positions (i.e., *d*_*i, j*_/*n*), is the maximum likelihood estimate of the probability of a recombination event with respect to a pair of haplotypes in a double haploid design (Wu et al. 2008a). It is also an estimate of the recombination fraction, so that these distances can be transformed into LOD scores (see Morton 1955). Missing data were dealt with in a pairwise manner, so that each pairwise comparison had missing data removed prior to estimation of *d*_*i, j*_/*n*. When values of *d*_*i, j*_/*n* exceeded 0.5, which is the theoretical maximum value given the expected 1:1 segregation pattern, they were set to 0.5. The *d*_*i, j*_/*n* values were used to construct the pairwise distance matrix between all possible pairs of loci passing our quality thresholds.

Loci were clustered hierarchically based on the pairwise distance matrix using Ward’s method as the linkage function (Ward 1963). The values of *d*_*i, j*_/*n* were squared prior to use of Ward’s method in hierarchical clustering. We explored groupings (*K*) based on clustering on the interval *K* = [8, 16]. This interval was chosen because it brackets the haploid chromosome number of foxtail pine (1*N* = 12). This entailed cutting the resulting dendrogram at a specific height, so that the desired number of groups resulted. Solutions were compared using silhouette widths for each locus (Rousseeuw 1987). The value of *K* which maximized the fraction of loci for which the silhouette width was maximal across the different values of *K* was selected as optimal.

Ordering of loci within clusters was carried out using MSTmap (Wu et al. 2008a). This method takes a full, undirected graph where nodes are loci and edges are based on the values of *d*_*i, j*_/*n* and finds the correct order of markers based on the minimum-weighted traveling salesman path (TSP). Wu et al. (2008a) showed that the minimum-weighted TSP can be found using a minimum spanning tree approach and that it corresponds to the correct order of the loci if the minimum spanning tree on the full, undirected graph is unique. We employed MSTmap using the maximum likelihood objective function, grouping turned off, imputation of missing data turned off, and the Kosambi mapping function (Kosambi 1944). The resulting ordering of loci within each cluster, along with the distances (i.e., cM) in each cluster, were taken as the initial linkage map from which data were error-corrected and imputed.

Data were subsequently imputed and corrected for errors using Maskov (Ward et al. 2013). A full account of the mechanics used in the algorithm of Maskov can be found in Text S1 from Ward et al. (2013). For our purposes, the accuracy of the imputation and error correction depends upon two choices: (1) the threshold for missing data for a given megagametophyte and (2) the number of contiguous loci where genotype errors can occur. We chose a value equal to 90% for the amount of missing data across megagametophytes for the former and a value of 5% of the number of loci in the initial map for each cluster for the latter (cf., Ward et al. 2013).

A final round of ordering was conducted with the imputed and error corrected data using MSTmap as described previously. Imputation and error correction resulted in many loci where *d*_*i, j*_/*n* = 0. These cosegregating markers were thus mapped to the same bin (Wu et al. 2008a). The collection of resulting ordered clusters was taken as the final linkage map for each of the four maternal trees. The end result of the linkage analysis was thus four independent linkage maps, one per maternal tree.

### Consensus Map Construction and Biological Interpretation

We took a two-step approach to the inference of the consensus linkage map. First, the four linkage maps, one for each maternal tree, were combined into a framework linkage map using MergeMap (Wu et al. 2008b). We constructed a set of weights with which to rank SNP orderings from each map as more or less probable based on the average amount of missing data, where a higher weight meant that the genotype data used to infer the linkage map had fewer instances of missing data (red: 0.05, green: 0.40, blue: 0.15, yellow: 0.40). Second, the remaining SNPs were added to the framework map by using the weighted average of the observed recombination fractions across libraries and constructing a linkage map as described previously based on these weighted average values. Consistency in the positioning and relative distances among framework markers was assessed using Spearman (Spearman 1904) and Mantel (Mantel 1967) correlations. Specifically, pairwise distances (cM) among framework markers were extracted from each linkage group on the framework map built using MergeMap as well as the map resulting from use of the weighted average recombination fractions in MSTmap. A Mantel correlation was used to test the null hypothesis that these distances were not correlated using a Bonferroni-corrected significance threshold of *α* = 0.05. Separate tests were performed for each of the 12 linkage groups. All analysis was conducted in the R ver. 3.0.2 statistical computing environment (R Core Team 2013).

Framework markers on the resulting consensus linkage map were used to estimate the size (Chakravarti et al. 1991) and coverage (Lange and Boehnke 1982) of the foxtail pine genome. The contigs from the assembly used to discover SNPs that appeared on the consensus linkage map were annotated using BLAST tools (Altschul et al. 1990) and the most recent release of the loblolly pine (*Pinus taeda* L.) genome sequence (v. 1.01, annotation V2). Each contig from the assembly was queried against the set of scaffolds comprising the loblolly pine genome using blastn. The hits from each comparison were retained and these top hits were filtered based on query coverage and the percent identity. As a thresholds, we used a minimum of 50% for the query coverage and 75% for the percent identity. The percent identity for the query coverage was set according to the expected number of substitutions between two sequences (2*µt*, see Nei 1987), where the mutation rate (*µ*) was assumed to be 1 *×* 10^−9^ substitutions/site/year and the divergence time (*t*) was assumed to be 8 × 10^7^ years (Willyard et al. 2007). This translated into an average expectation of 16% divergence between any two DNA sequences of loblolly and foxtail pines. We rounded down to 75% to account for a portion of the variance around this expectation. Hits that exceeded these thresholds were transferred as annotations, as obtained from the annotation gff files, to the contig appearing on the consensus linkage map for foxtail pine. The resulting GO annotations were visualized and analyzed with ReviGO (Gene Ontology monthly release 10/2014; UniProt-to-GO mapping 9/30/2014) (Supek et al. 2011) allowing a similarity of 50% across terms.

## Results

### DNA Sequence Analysis

The raw number of reads varied across libraries from a minimum of 71 834 280 (red) to a maximum of 206 365 836 (green), with an average of 153 082 376 ± 49 855 941. All raw reads were either 102 bp (green, yellow) or 110 (red, blue) in length, depending on sequencing technology (HiSeq 2500 vs 2000, respectively). In general, the libraries run on the Illumina HiSeq 2500 platform had a 1.65-fold greater number of reads than those run on the Illumina 2000 HiSeq platform. Processing of reads for quality reduced these numbers by approximately 1.66-fold, with a range of a 2.56-fold (red) to a 1.33-fold (yellow) reduction. After filtering, the average length of reads was 88 ± 13 bp, with a range of 40 bp to 102 bp across libraries. The number of quality-filtered reads per megagametophyte also varied 19 741-fold (± 27 069-fold) on average across libraries, with average minimums of 753 ± 603 bp to average maximums of 3 421 571 ± 2 070 990 bp. After quality filtering this translated into an average total of 8 137 663 036 bp ± 3 658 147 958 bp generated per library, ranging from 2 436 531 265 bp (red) to 11 643 165 529 bp (yellow).

The largest number of reads (*n* = 6 838 986) were obtained for a single megagametophyte in the green library. These reads were used to create an assembly against which all other data were mapped for SNP calling and genotype determination. Optimization of assembly parameters (*k* = 31, *lnL* = *−*110.071), resulted in an assembly of 231 053 contigs, with an average length of 89 bp ± 12 bp per contig (range: 61 bp to 312 bp), and an average per-contig base coverage of 4.5 X to 20.0 X (range: 1.5 X to 5069 X). This assembly represented approximately 0.07% of the genome of foxtail pine, which was assumed to be approximately 30 Gbp in size (Murray 1998).

**Table 1.** Attributes of the data structure related to maternal tree.

| Attribute | Yellow | Blue | Red | Green |
| --- | --- | --- | --- | --- |
| Region | Klamath | Klamath | Sierra Nevada | Sierra Nevada |
| Latitude | 44.7483 | 41.1959 | 36.4481 | 36.4481 |
| Longitude | -123.1332 | -122.7922 | -118.1706 | -118.1706 |
| Illumina Platform | HiSeq 2500 | HiSeq 2000 | HiSeq 2000 | HiSeq 2500 |
| # Megagametophytes | 95 | 95 | 76 | 73 |
| Reads (total) | 174 516 834 | 159 612 555 | 71 834 280 | 206 365 836 |
| Reads (filtered) | 131 540 433 | 80 498 688 | 28 041 978 | 129 396 774 |

**Fig. 2.**
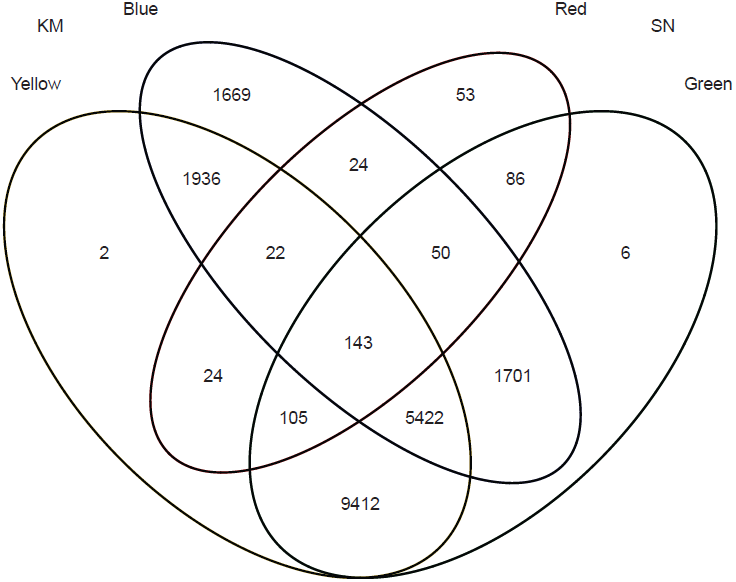
Sharing of contigs across maternal tree maps from which the consensus map was constructed. Counts in each cell represent the number of unique contigs appearing on the final consensus map. Unique contigs for the yellow and green maternal trees were largely discarded to make estimation of pairwise recombination fractions computationally feasible (see Materials and Methods). KM = Klamath Mountains; SN = Sierra Nevada.

Using this assembly, 349 542 putative SNPs were called (Table 1). These 349 542 SNPs were located in 83 051 unique contigs (35.94% of the total), with a mean of 4 SNPs per contig (range: 1 to 32). Filtering these SNPs by expected segregation patterns, consistency with heterozygous calls for the psuedo-diploid sample, and minimum sample sizes for genotype calls, resulted in 983, 34 261, 21 594, and 35 304 SNPs for the red, green, blue, and yellow libraries, respectively. The vast majority of SNPs eliminated were for violation of the 1:1 expected pattern of segregation (259 801 - 268 621), with approximately 95% of these dropped SNPs shared across families. The counts for the yellow and green libraries were also trimmed so as to remove all but a handful (*n* = 2 for the yellow and *n* = 6 for green libraries) of the unique contigs not found as polymorphic in the other libraries. This was done to facilitate the efficiency of the calculation of pairwise recombination fractions. These SNP counts represented 507, 16 925, 10 967, and 17 066 contigs for the red, green, blue, and yellow libraries, respectively. Patterns of shared polymorphic contigs, as well as SNPs, were as expected given the among-region magnitude of genetic differentiation (Figure 2, see Eckert et al. 2008), with libraries comprised of megagametophytes sampled from maternal trees located in the same geographical area sharing more polymorphic contigs and SNPs than comparisons of maternal trees from different geographical regions (nonparametric permutation analysis: *P <* 0.0001, see Supplemental Text). On average, megagametophytes in the filtered data set had 79.40% (± 14.7%) missing data (i.e., a missing haploid genotype) across SNPs (range: 1.3% to 99.8%), with the green library having the smallest (74.3% ± 18.9%) and the red library having the largest average amount of missing data per megagametophyte (84.4% ± 15.2%).

### Linkage Mapping

Individual linkage maps were constructed for each maternal tree separately using an iterative approach based on imputation. All filtered SNPs for each maternal tree, regardless of being located in the same contig, were assessed for patterns of linkage followed by grouping and ordering of SNPs. Redundant SNPs were filtered *post hoc* and used to test for biases in our analysis pipeline.

Grouping of pairwise recombination fractions via hierarchical clustering was consistent with 12 linkage groups. This corresponded to a minimum pairwise LOD score of approximately 5.5 for each maternal tree for markers to be placed within the same linkage group. Inspection of the distribution of silhouette values for values of *K* ranging from 8 to 16 revealed that *K* = 12 was the best clustering solution for each of the 4 maternal trees (Figure S1). This was confirmed by comparison of pairwise LOD scores for SNPs within versus among the 12 linkage groups. Comparisons within linkage groups were on average 3.2-fold larger than among linkage groups, which was significantly greater than expected randomly (*n* = 1, 000 permutations/maternal tree, *P* < 0.015).

Marker ordering within putative linkage groups using MSTmap resulted in extremely long linkage maps (e.g., *>* 50 000 cM) for each maternal tree. This translated into an average number of recombination events which exceeded 100 per megagametophyte. This pattern is consistent with problems of inference due to missing data and genotyping errors (Ward et al. 2013). To verify this assumption, data for the blue library were split into two sets of 35 megagametophytes - those with the least amount of missing data and those with the largest amount of missing data. As expected, the inferred recombination distances were 3.5-fold smaller for the maps inferred using the megagametophytes with less missing data. Thus, we followed the approach of Ward et al. (2013) to impute and error correct data based on our initial marker orderings.

Imputation and error correction of genotype data for each linkage group for each maternal tree was carried out using Maskov. This process drastically reduced the number of inferred recombination events, including double crossovers, from *>* 100 per megagametophyte to approximately 1 to 2 per megagametophyte. This reduction was controlled by setting a parameter in Maskov so as to produce a number of recombination events per megagametophyte that mirrored those observed previously for linkage mapping within conifers (Eckert et al. 2009; Martinez-Garcia et al. 2013). Changing this parameter had no effect on the downstream ordering of SNPs within linkage groups, but only changed the spatial resolution of the resulting linkage map.

The resulting linkage maps for each maternal tree were aligned manually based on the presence of shared contigs. Overall, there was excellent agreement among maps, with only 115 SNPs being mapped to conflicting linkage groups across maternal trees. All 115 SNPs with conflicting group assignments were unique to the red library. These were dropped from further consideration. Within linkage groups, SNPs present in multiple libraries were ordered similarly (pairwise Spearman’s *ρ >* 0.956, *P* < 0.001), with conflicting orderings having average differences of 5.91 cM (±5.64 cM). Inferred linkage maps for each maternal tree also resulted in SNPs from the same contig largely being mapped to the same position, with an average of only 5.8% of SNPs from the same contig being mapped to a different position. Approximately 94% of the time, these different positions were adjacent on the linkage map. For those SNPs from the same contig that did not map to the same position, the average difference in positioning was 1.64 cM (± 3.01 cM), with no instances of SNPs from the same contig being located on different linkage groups. We thus pruned multiple SNPs per contig by randomly selecting one SNP per contig from the data set and re-estimated the linkage maps for each maternal tree as described previously. The resulting 4 linkage maps were taken as the final estimates of linkage relationships among polymorphic contigs in each of the 4 maternal trees.

**Table 2.**
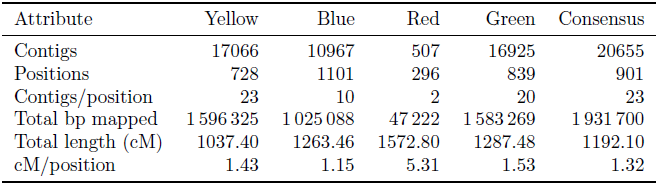
Attributes of single-tree and the consensus linkage maps. Values for ratio variables are totals and are not averaged across linkage groups (see Tables S1–S5).

| Attribute | Yellow | Blue | Red | Green | Consensus |
| --- | --- | --- | --- | --- | --- |
| Contigs | 17066 | 10967 | 507 | 16925 | 20655 |
| Positions | 728 | 1101 | 296 | 839 | 901 |
| Contigs/position | 23 | 10 | 2 | 20 | 23 |
| Total bp mapped | 1 596 325 | 1 025 088 | 47 222 | 1 583 269 | 1 931 700 |
| Total length (cM) | 1037.40 | 1263.46 | 1572.80 | 1287.48 | 1192.10 |
| cM/position | 1.43 | 1.15 | 5.31 | 1.53 | 1.32 |

**Fig. 3.**
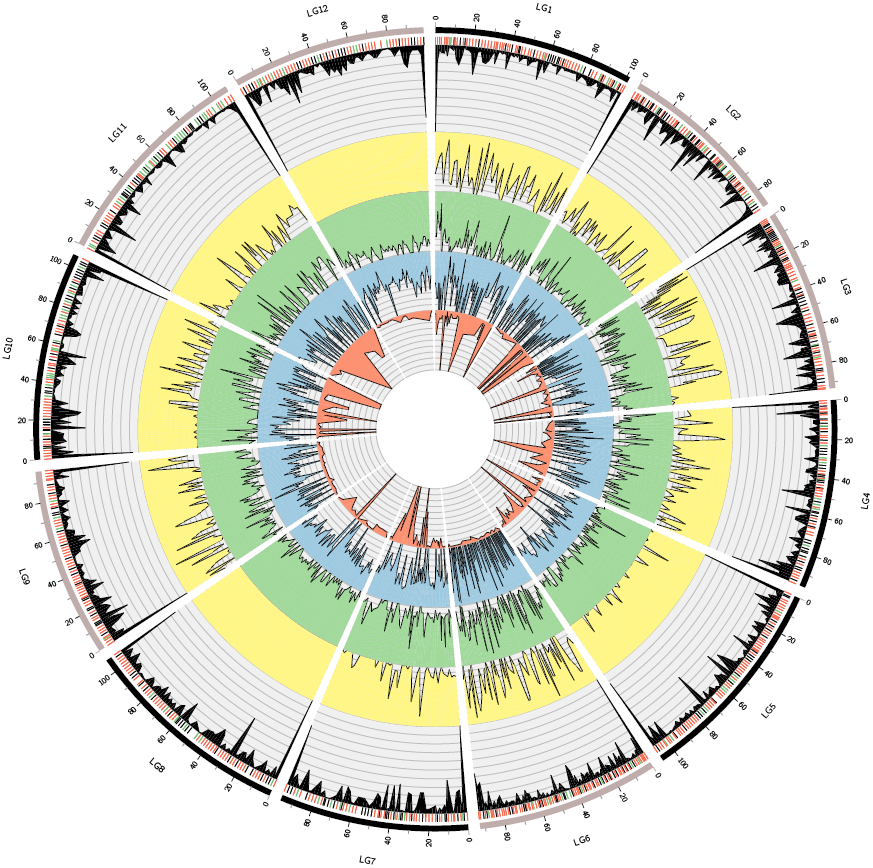
Consensus linkage map of 12 linkage groups, derived from SNPs among individuals of four populations. Working inward from the outermost section of the figure, for each linkage group: (1) the solid black bars represent the span of recombination distances (in centiMorgan) for markers; (2) the individual tick marks show the locations of the markers and the colors represent the density of annotation of the SNPs at that position ((*≥* 50% = green, *≥* 25%= red, *<* 25% = black) to homologous locations in lobolly pine); (3) The black density plot represents the counts of SNPs from all four families mapping to a specific position in the linkage group; (4) the colored density plots show the contribution SNPs from the individual families to the markers on the map at each position, and are shown in order by total read count in the library, with yellow having the most and red having the least amount of reads. Linear plots of linkage groups comprising the consensus map are given in Figure S9.

The final 4 linkage maps varied in total length from 1037.40 cM to 1572.80 cM, with an average of 1290.29 cM (± 219.5 cM; Tables 2, S1–S4; Figures 3, S2–S8). In total, 20 655 unique contigs representing 1 931 700 bp of DNA were mapped to a position within at least one linkage map. The number of contigs varied 33.66-fold across linkage maps, with a minimum of 507 (red) to a maximum of 17 066 (yellow). These contigs were organized into an average of 741 (± 335) unique positions, separated on average by 1.77 cM (± 2.36 cM), across linkage maps, with the fewest number of unique positions observed in the linkage map for the red maternal tree (*n* = 296) and the largest number in the linkage map for the blue maternal tree (*n* = 1101). With respect to average distances between adjacent positions, the linkage map for the red maternal tree had the largest (5.53 cM ± 6.11 cM), while that for the blue maternal tree had the lowest (1.16 cM ± 0.77 cM). This translated into an average of 15 (± 27) contigs per position on average, with the linkage map for the red maternal tree having the fewest contigs per position on average (2 ± 2) and the linkage map for the yellow maternal tree the most contigs per position on average (27 ± 35). Contigs were also non-randomly distributed across positions for all linkage maps except that for the red maternal tree (*P <* 0.0001, see Supplemental Text), with elevated contig counts typically occurring at the ends of linkage groups (Figure 3).

### Consensus Map Construction and Biological Interpretation

A set of 507 framework SNPs were devised from those contigs shared across at least three of the four linkage maps. These 507 SNPs were used to construct a framework map using MergeMap. The resulting linkage map had an overall length of 1572.80 cM. Comparison of this map with those for each maternal tree revealed a strong similarity in positioning for each linkage group (Spearman’s *ρ >* 0.98, *P <* 0.001). Using an expanded set of SNPs present in at least three families, two of which had to be the green and yellow families, confirmed these patterns, with pairwise correlations among maps on the order of 0.92 or greater. Given this overall similarity, we incorporated the remaining markers into the map by using weighted averages of observed pairwise recombination fractions across maternal trees and inferred a consensus linkage map as outlined previously. Inferred marker positions and distances for the framework markers were highly correlated across linkage groups in this map relative to that inferred using MergeMap and only the framework markers (Mantel’s *r* :*>* 0.95*, P <* 0.001). We used this as evidence in support of our approach and the inferred consensus linkage map was taken as the final consensus estimate of linkage relationships for the 20 655 unique contigs located in the four maternal tree linkage maps.

As with the individual tree maps, *K* = 12 linkage groups was most consistent with the averaged data. This corresponded to a minimum pairwise LOD score of approximately 5.5 for each maternal tree for markers to be placed within the same linkage group. The consensus linkage map was 1192.00 cM in length, with linkage groups varying in length from 88.44 cM to 108.76 cM (Table S5, Figures 3, S9). There were 901 unique positions across the 12 linkage groups for this map, so that the average number of contigs per position was 23 (± 35). These 901 positions were separated on average by 1.34 cM (± 0.50 cM). As with the individual maternal tree linkage maps, contigs were non-randomly distributed across positions (*P <* 0.0001, see Supplemental Text), with notable enrichment at the ends of inferred linkage groups. Using the 507 framework SNPs and the final consensus linkage map, the estimated genome size of foxtail pine is 1276.04 cM (95% confidence interval: 1174.31 - 1377.77 cM). As such, the estimated coverage of the genome is 98.58% (LOD threshold = 5.5, maximum distance among adjacent framework markers: 13.4 cM, number of framework markers: 507, *K* = 2694).

Of the 20 655 contigs in the reference assembly which contained SNPs, 5627 (27.2%) contained BLASTn hits (*n* = 5853) which passed the filtering threshold of 50% query length and 75% identity. The averages of query length (bp), query coverage percentage, and identity percentage were 94 ± 7, 86.4% ± 13.0%, and 90.1% ± 3.9%, respectively. We found 2802 (48.9 %) instances of SNPs mapping to putative genic regions in the loblolly pine genome (*n* = 587 scaffolds) representing 303 unique GO terms. More detailed annotation information (e.g., InterPro IDs and GO terms) can be found in the supplemental file S1.

### Discussion

The genetic architecture of fitness-related traits has been a major focus of geneticists for over a century (reviewed by Ellegren and Sheldon 2008). Early efforts to understand the genetic architecture of fitness-related traits focused primarily on the number and effect size of the loci underlying heritable, phenotypic variation (Fisher 1918). Recent work has extended this line of research, with a multitude of studies linking phenotypic with genetic variation through linkage mapping, both within pedigrees (Mauricio 2001; Neale and Kremer 2011; Ritland et al. 2011) and within natural populations (Ingvarsson and Street 2011; Eckert et al. 2013), or through quantitative genetic experimentation (Anderson et al. 2014, 2013; Fournier-Level et al. 2013). Relatively little empirical work outside of model organisms, other than polyploidization or the characterization of genomic islands of divergence (e.g., high F_ST_ ), has focused on the genomic organization of loci contributing to fitness differences among individuals (but see Stevison et al. 2011). This is despite clear theoretical predictions relating the evolution of the genetic architecture underlying fitness-related traits to the genomic organization of the loci comprising this architecture (Kirkpatrick and Barton 2006; Yeaman and Whitlock 2011; Yeaman 2013; Akerman and Burger 2014).

Here, we have provided a high-density linkage map representing over 20,000 unique contigs distributed throughout the 30 Gbp genome of foxtail pine that can be used to aid in the discovery and study of loci contributing to local adaptation. To our knowledge it represents one of the most dense linkage maps ever produced within forest trees, although the number of unique positions is much less than the number of mapped contigs (i.e., about 1/20^*th*^). Approximately 25% of these contigs had significant similarity to sequences within the draft loblolly pine genome. Importantly, our markers are dispersed in both genic and non-genic regions of the genome. The latter are often ignored in studies of local adaptation utilizing markers based on sequence capture (e.g., Neves et al. 2014) and SNP arrays (e.g., Eckert et al. 2010b), yet it is known that non-genic regions are often involved with adaptation (e.g., Studer et al. 2011). This linkage map, moreover, was created using affordable next generation sequencing technologies in combination with freely-available methods of analysis, which highlights the feasibility of this approach to non-model conifers, where full genome sequencing and assembly are still not quite feasible given realistic research budgets. With regard to map integrity, recombination fractions for pairs of SNPs segregating in multiple trees were highly correlated (Mantel’s *r* > 0.90 for all comparisons across linkage groups). This allowed for the creation of a robust consensus linkage map, as well as highlighted the biological signal of linkage apparent even in noisy ddRADseq data. When coupled with the other biological signals in our results (e.g., trees from the same regional population sharing SNPs more often), we can be confident that our inferred linkage maps are based primarily on biological, as opposed to statistical, signals. In further support of this claim, randomly subsampling our data to represent 10 000 contigs and performing linkage mapping as described previously resulted in a consensus linkage map that was indistinguishable from that pictured in Figure 3 (Spearman’s *ρ* = 0.997*, P <* 0.0001).

The linkage map produced here is valuable in numerous ways. First, it provides a dense resource for quantitative trait locus (QTL) mapping. Our next step using this linkage map is to link fitness-related phenotypic variation with genotypes at mapped markers. We are currently mapping QTLs for *δ*^13^C to accomplish this goal (cf., Hausmann et al. 2005). Importantly, this will represent one of the first QTL maps in the clade of soft pines outside of section *Quinquefoliae*. Second, the framework provided here is optimal for imputation and phasing of data during population genomic inferences utilizing samples from natural populations (Scheet and Stephens 2006). Third, our linkage map is the foundation upon which theoretical expectations can be tested. For example, the theory of Yeaman and Whitlock (2011)predicts that the loci contributing to local adaptation should be differentially clustered in genomes as a function of rates of gene flow among populations. Magnitudes of gene flow among stands differ dramatically within the regional populations of foxtail pine, and pairwise plots of synteny across maternal trees revealed several instances of differential marker orderings between regional populations consistent with areas of the genome with structural differences (see Figures S7-S8). These areas, however, were the exception, as marker orderings across trees were highly correlated. Fourth, knowledge about the physical ordering of loci allows patterns of linkage disequilibrium (LD) within natural populations to be better characterized. The role of LD in local adaptation has long been recognized (see Akerman and Burger 2014), yet empirical studies of its role are difficult without some knowledge of physical relationships among loci. This is because LD among physically linked markers is expected to some degree, whereas LD among physically unlinked markers must have originated from some evolutionary process (e.g., genetic drift, natural selection, migration). Patterns of LD across non-genic regions of pine genomes are currently unknown (but see Moritsuka et al. 2012), so additional data in combination with the linkage map provided here allow for rigorous investigations of these patterns. Lastly, continued production of linkage maps across the Pinaceae will aid comparative genomics and evolutionary inference through study of synteny and the evolution of genome structure (Ritland et al. 2011; Pavy et al. 2012).

Despite numerous indicators of biological signals dominating our dataset, caution is still needed when interpreting our results. First, we used novel analysis methods that have not been tested using simulations. For example, the form of hierarchical clustering used here is not employed to our knowledge in any of the available software packages used for linkage mapping. Its utility on data of smaller or larger sizes than that presented here is unknown. Consistency of results across maternal trees, however, indicates that our methods are likely appropriate for our data (see also Tani et al. 2003). Second, error-correction and imputation were used, which could have affected marker ordering and distances. Marker order, however, did not change with increasing stringency of error correction. Only marker distances changed with increased stringency, thus creating clumped distributions of makers. This was also apparent in the total map length, which is at the lower end expected for conifers (cf., Ritland et al. 2011), which is indicative of being conservative with error corrections. The effect of marker clumping on downstream uses of this linkage map, however, is likely to be minimal (e.g. bias in QTL intervals), as this bias would affect QTL size and not necessarily inference of QTL presence or absence. The relative importance of imputation and error correction is to some degree affected by experimental conditions. We did not standardize the total amount of DNA for each megagametophyte prior to construction of libraries (concentration ranges: 10 ng/ul to >50 ng/ul, which likely affected the average 19 741-fold variation in the number of reads across megagametophytes. Future studies would benefit from considering this prior to library construction. Related to this issue was the poor performance of the red family. In 506 general, the library for this family exhibited signs of low sequence quality, with it having the largest fraction of reads eliminated during quality filtering (Table 1), the largest fraction of pseudo-diploid genotypes called as homozygous or missing (99.5%), and the largest fraction of loci deviating from expected segregation patterns (76.8%). This is consistent with lower overall coverage driven by low quality sequence data, which could have resulted from any of the numerous laboratory steps during creation of the multiplexed libraries (i.e., DNA extractions, restriction digests, ligation, and PCR). Third, we used a form of hierarchical clustering that required the number of groups to be defined subjectively. *Post hoc* analysis indicated that our clustering solution corresponded to a LOD threshold of approximately 5.5 and that 12 was an optimal number of groups (Figure S1). Selection of a larger or smaller number of groups, moreover, did not change marker ordering within groups substantially. Typically, changing *K* to a larger value broke existing linkage groups into more pieces, whereas changing *K* to smaller values merged existing linkage groups. Marker orderings within these broken or merged groups, however, did not change. Fourth, we did not explicitly quantify error rates. In theory, error rates can be calculated from summaries derived from the mpileup in samtools. Given the relatively low coverage and limited reference assembly for this species (cf., Nystedt et al. 2013; Neale et al. 2014), estimation of error rates would likely be biased. Thus, we preferred to acknowledge the presence of errors, as indicated by the extremely long initial single-tree linkage maps, and use statistical methods to minimize their influence. As next generation data accumulate for this species, and other conifers in general, precise estimation of error rates will become feasible. Lastly, our sample sizes were not large enough to resolve linkage relationships beyond distances of approximately 1.0 cM (but see Neves et al. 2014). Increased number of sampled megagametophytes would have allowed higher resolution, which could aid in downstream uses of our linkage map. Despite this level of resolution, however, we have produced one of the densest linkage maps to date for a forest tree species (Eckert et al. 2010a; Martinez-Garcia et al. 2013; Neves et al. 2014).

Conifer genomics is emerging as a mature scientific field (Mackay et al. 2012). Draft sequences of genomes and transcriptomes for several species have been released and more are planned. As shown here, production of high-density linkage maps is a fruitful endeavor to accompany this maturation. The results presented here are promising and also provide guidance for future attempts in additional species. Specifically, linkage maps provide ample information about genomic structure that is needed for the study of local adaptation in natural populations (cf., Limborg et al. 2014). Here, we have produced a high-density linkage map for foxtail pine using methods applicable to any non-model conifer species, thus opening the door for further studies of genome structure and the genetic architecture of local adaptation in this rather understudied clade of pines, as well as the Pinaceae as a whole.

## Acknowledgements

The authors would like to thank the staff at the USDA Institute of Forest Genetics, the VCU Nucleic Acids Research Facility, and the VCU Center for High Performance Computing. In addition, we would like to thank Tom Blush and Tom Burt for help in obtaining seeds. Funding for this project was made available to AJE via start-up funds from Virginia Commonwealth University. CJF was supported by the National Science Foundation (NSF) National Plant Genome Initiative (NPGI): Postdoctoral Research Fellowship in Biology (PRFB) FY 2013 Award #NSF-NPGI-PRFB-1306622.

## Data Archiving Statement

Raw short read data are located in the NCBI Short Read Archive (accession number: PRJNA266319). The linkage map summary files, assembly used for read mapping and SNP calling, and VCF files are given as supporting documents (Files S1 - S3). The consensus linkage map is also available in the Comparative Mapping Database located at the Dendrome website (accession number: TG151). Source code for this manuscript and data analyses are located at http://www.github.com/cfriedline/foxtail_linkage.

## Supplementary Materials

### Supplementary Text

A nonparametric permutation analysis was used to test the hypothesis that sharing of polymorphic contigs was greater between maternal trees located in the same regional population. The null hypothesis in this case is that the degree of sharing between trees in the same regional population is not different than between trees in different regional populations. To conduct this test, we constructed a null distribution of the difference between mean within versus mean between levels of contig sharing. This distribution was based on permutations, where each permutation consisted of randomly sampling with replacement contigs from the full set of contigs prior to filtering for each maternal tree identifier. The number of each contigs for each maternal tree identifier was equal to that observed in the original dataset. Given these assignments, the degree of sharing (DS) between trees was calculated as:

DS = number of contigs shared/number of unique contigs in the pairwise comparison

Given four maternal trees, there are 6 pairwise comparisons, of which there are 2 within regional population and 4 between regional population comparisons. Using these pairwise values, we constructed a test statistic defined as the difference between the mean within regional population value of DS and the mean between regional population value of DS. We simulated 10 000 values of this test statistic to form the null distribution and rejected the null hypothesis when the observed test statistic fell in the upper 95% tail of this null distribution (i.e., a one-tailed test). Using this approach, the observed test statistic was 0.06337814. The limits of the simulated null distribution were -0.0349 to 0.0415. Thus the observed test statistic fell completely outside the upper tail of the null distribution, which gives *P <* 0.0001.

The spatial distribution of contigs along linkage groups was tested against complete spatial randomness using simulations. Specifically, we used the variance in the observed number of contigs mapped to each unique position across the entire linkage map as the test statistic. The null distribution of the test statistic was created by simulating 10 000 Poisson-distributed variables, each with the number of random values equal to the number of unique positions on the linkage map under consideration and the mean equal to the mean number of contigs per position. For each Poisson-distributed variable, the variance was calculated. The set of 10000 variances was used as the null distribution expected under complete spatial randomness. A separate null distribution was constructed for each maternal tree. If the observed variance fell in either the 2.5% or 97.5% tail of the null distribution, the null hypothesis was rejected. For example, the values used for the consensus linkage map were: observed variance = 1245.423, average expected variance of Poisson-distributed variable = 23 with 95% CI of null expectation of 20.915–25.246, and limits of the null distribution of 19.214– 27.655. The *P* - value for the observed variance is thus *P <* 0.0001.

This test does not explicitly test spatial randomness of the unique positions on a linkage map, but rather tests the assumption that the intensities (i.e., the number of mapped contigs) at each position follow a Poisson distribution. This approach was chosen because distances in the inferred linkage maps were sensitive to the error correction and imputation. Of course, the number of contigs at each mapped position is also sensitive to this process, although relaxing the error correction parameter in Maskov resulted in similar results for a range of values that resulted in realistic linkage map lengths (i.e., total lengths < 3000 cM). Thus, we were confident that this result is not only a function of error correction and imputation. All linkage maps except for the single-tree map from the red maternal tree had greater than expected variances across positions in counts of mapped contigs relative to the null model of a Poisson-distributed variable.

**Table S1.** Attributes of the single-tree linkage by linkage group (LG) for the yellow maternal tree (Klamath).

| Attribute | No. contigs | No. positions | Contigs/position | Total bp mapped | Total length (cM) | cM/position |
| --- | --- | --- | --- | --- | --- | --- |
| LG1 | 1394 | 59 | 24 | 130 466 | 85.17 | 1.44 |
| LG2 | 1518 | 52 | 29 | 142 195 | 71.07 | 1.37 |
| LG3 | 1174 | 50 | 23 | 110 032 | 69.35 | 1.39 |
| LG4 | 879 | 58 | 15 | 82 151 | 86.87 | 1.50 |
| LG5 | 2046 | 79 | 26 | 191 663 | 109.62 | 1.39 |
| LG6 | 1893 | 51 | 37 | 176 401 | 67.97 | 1.33 |
| LG7 | 1681 | 56 | 30 | 157 609 | 79.47 | 1.42 |
| LG8 | 1863 | 70 | 27 | 173 888 | 97.32 | 1.39 |
| LG9 | 1480 | 63 | 23 | 138 205 | 86.86 | 1.38 |
| LG10 | 872 | 68 | 13 | 81 645 | 97.32 | 1.43 |
| LG11 | 578 | 61 | 9 | 54 056 | 101.42 | 1.66 |
| LG12 | 1688 | 61 | 28 | 158 014 | 84.96 | 1.39 |

**Table S2.** Attributes of the single-tree linkage by linkage group (LG) for the blue maternal tree (Klamath).

| Attribute | No. contigs | No. positions | Contigs/position | Total bp mapped | Total length (cM) | cM/position |
| --- | --- | --- | --- | --- | --- | --- |
| LG1 | 1085 | 128 | 8 | 101 420 | 112.69 | 0.88 |
| LG2 | 982 | 81 | 12 | 91 963 | 102.56 | 1.27 |
| LG3 | 1306 | 113 | 12 | 121 886 | 103.84 | 0.92 |
| LG4 | 610 | 85 | 7 | 57 028 | 96.18 | 1.13 |
| LG5 | 904 | 76 | 12 | 84 711 | 108.75 | 1.43 |
| LG6 | 1769 | 129 | 14 | 165 198 | 108.89 | 0.84 |
| LG7 | 879 | 89 | 10 | 82 216 | 101.30 | 1.14 |
| LG8 | 805 | 66 | 12 | 75 042 | 108.89 | 1.65 |
| LG9 | 1001 | 98 | 10 | 93 821 | 98.76 | 1.01 |
| LG10 | 470 | 93 | 5 | 44 024 | 108.90 | 1.17 |
| LG11 | 365 | 84 | 4 | 34 052 | 111.42 | 1.33 |
| LG12 | 791 | 59 | 13 | 73 727 | 101.28 | 1.72 |

**Table S3.** Attributes of the single-tree linkage by linkage group (LG) for the red maternal tree (Sierra Nevada).

| Attribute | No. contigs | No. positions | Contigs/position | Total bp mapped | Total length (cM) | cM/position |
| --- | --- | --- | --- | --- | --- | --- |
| LG1 | 100 | 26 | 4 | 9388 | 123.44 | 4.75 |
| LG2 | 24 | 16 | 2 | 2205 | 47.54 | 2.97 |
| LG3 | 50 | 34 | 1 | 4642 | 134.27 | 3.95 |
| LG4 | 34 | 18 | 2 | 3202 | 133.33 | 7.41 |
| LG5 | 50 | 40 | 1 | 4631 | 169.48 | 4.24 |
| LG6 | 35 | 19 | 2 | 3191 | 136.74 | 7.20 |
| LG7 | 65 | 43 | 2 | 6059 | 135.49 | 3.15 |
| LG8 | 30 | 18 | 2 | 2814 | 125.48 | 6.97 |
| LG9 | 37 | 22 | 2 | 3428 | 143.35 | 6.52 |
| LG10 | 38 | 33 | 1 | 3511 | 166.06 | 5.03 |
| LG11 | 16 | 10 | 2 | 1501 | 108.08 | 10.81 |
| LG12 | 28 | 17 | 2 | 2650 | 149.54 | 8.80 |

**Table S4.**
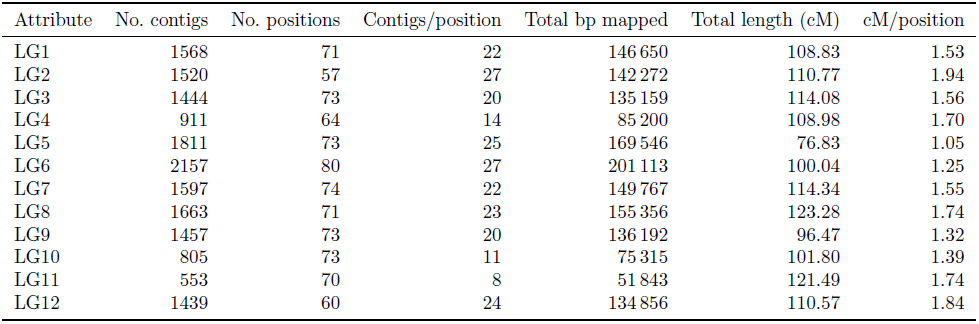
Attributes of the single-tree linkage by linkage group (LG) for the green maternal tree (Sierra Nevada).

| Attribute | No. contigs | No. positions | Contigs/position | Total bp mapped | Total length (cM) | cM/position |
| --- | --- | --- | --- | --- | --- | --- |
| LG1 | 1568 | 71 | 22 | 146 650 | 108.83 | 1.53 |
| LG2 | 1520 | 57 | 27 | 142 272 | 110.77 | 1.94 |
| LG3 | 1444 | 73 | 20 | 135 159 | 114.08 | 1.56 |
| LG4 | 911 | 64 | 14 | 85 200 | 108.98 | 1.70 |
| LG5 | 1811 | 73 | 25 | 169 546 | 76.83 | 1.05 |
| LG6 | 2157 | 80 | 27 | 201 113 | 100.04 | 1.25 |
| LG7 | 1597 | 74 | 22 | 149 767 | 114.34 | 1.55 |
| LG8 | 1663 | 71 | 23 | 155 356 | 123.28 | 1.74 |
| LG9 | 1457 | 73 | 20 | 136 192 | 96.47 | 1.32 |
| LG10 | 805 | 73 | 11 | 75 315 | 101.80 | 1.39 |
| LG11 | 553 | 70 | 8 | 51 843 | 121.49 | 1.74 |
| LG12 | 1439 | 60 | 24 | 134 856 | 110.57 | 1.84 |

**Table S5.** Attributes of the consensus linkage by linkage group (LG).

| Attribute | No. contigs | No. positions | Contigs/position | Total bp mapped | Total length (cM) | cM/position |
| --- | --- | --- | --- | --- | --- | --- |
| LG1 | 1938 | 80 | 24 | 181 309 | 101.64 | 1.27 |
| LG2 | 1837 | 73 | 25 | 172 031 | 88.44 | 1.21 |
| LG3 | 1977 | 77 | 26 | 184 680 | 93.10 | 1.21 |
| LG4 | 1108 | 73 | 15 | 103 585 | 96.23 | 1.32 |
| LG5 | 2068 | 82 | 25 | 193 722 | 107.35 | 1.31 |
| LG6 | 2843 | 96 | 30 | 265 356 | 92.59 | 0.96 |
| LG7 | 1871 | 70 | 27 | 175 272 | 96.22 | 1.37 |
| LG8 | 1863 | 70 | 27 | 173 888 | 107.16 | 1.53 |
| LG9 | 1818 | 71 | 26 | 169 974 | 95.89 | 1.35 |
| LG10 | 956 | 76 | 13 | 89 477 | 106.06 | 1.40 |
| LG11 | 687 | 71 | 10 | 64 299 | 108.76 | 1.53 |
| LG12 | 1689 | 62 | 27 | 158 107 | 98.56 | 1.60 |

**Fig. S1.**
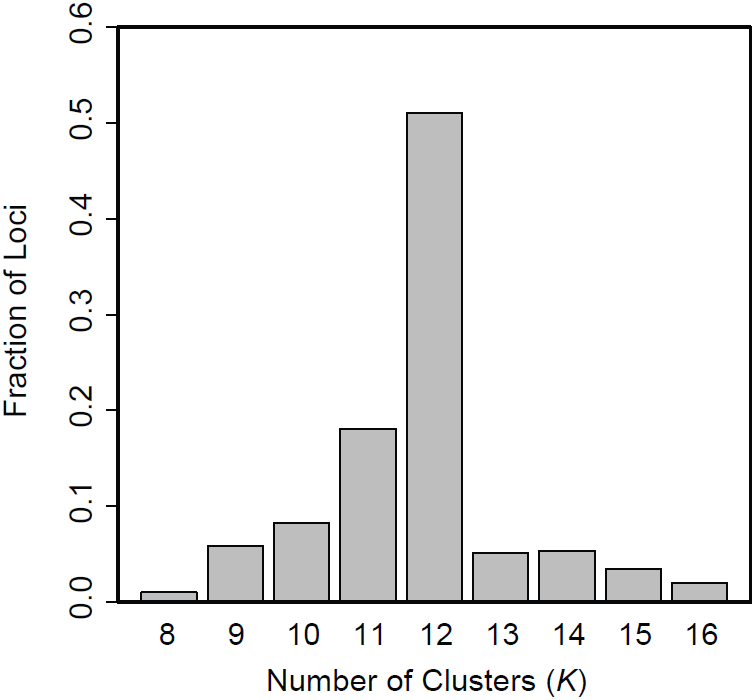
The fraction of loci with silhouette values being maximal at each value of *K* reveals that *K* = 12 is an optimal clustering solution. For each locus, the maximum silhouette value was determined and the fraction of loci with maximal values at each value of *K* was plotted. These results are for the yellow maternal tree. Results for the other single-tree, as well as consensus linkage map were qualitatively similar with pronounced peaks at *K* = 12.

**Fig. S2.**
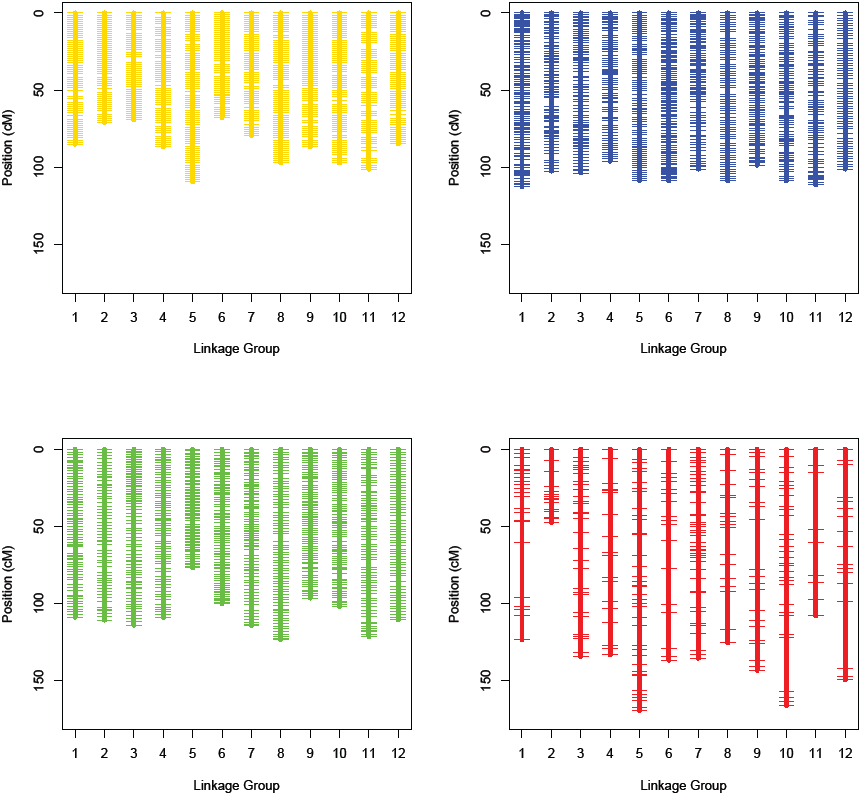
Single-tree linkage maps. Maps are color coded and organized by regional population (top row: Klamath; bottom row: southern Sierra Nevada

**Fig. S3.**
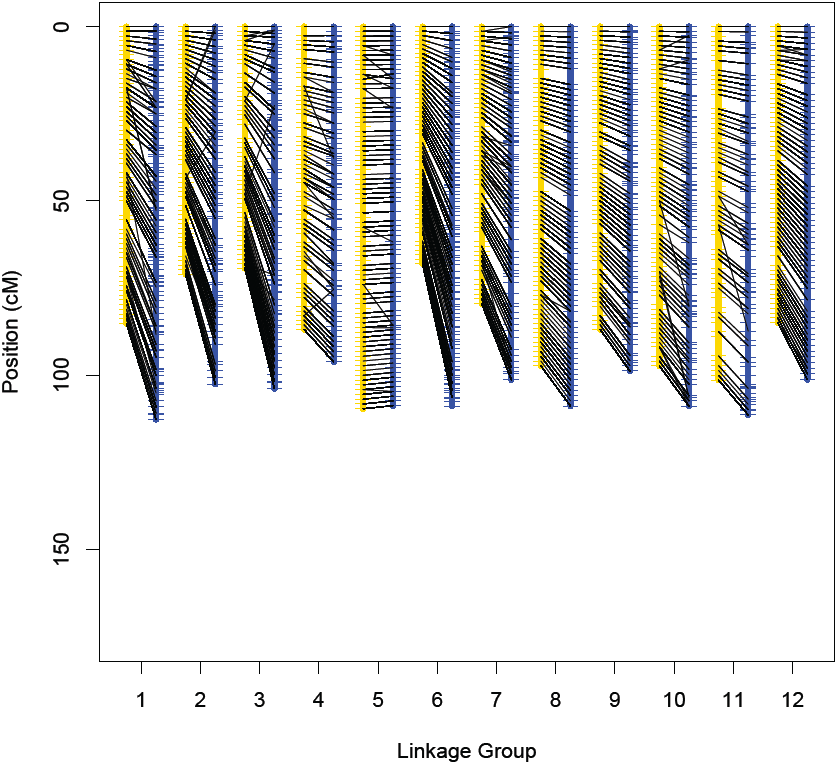
Pairwise synteny plots between the yellow and blue maternal trees. Both trees are from the Klamath region.

**Fig. S4.**
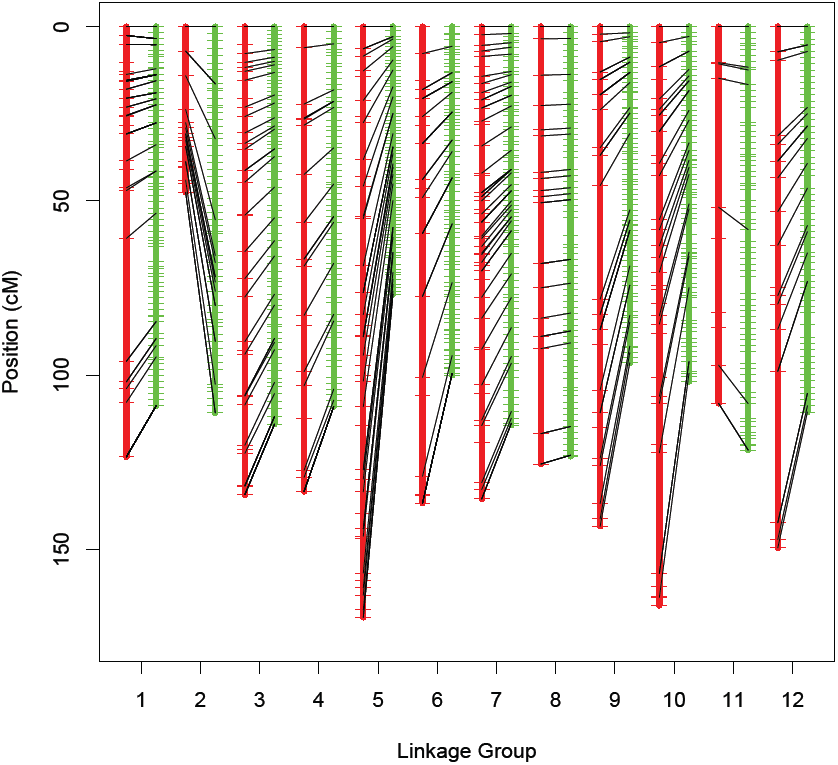
Pairwise synteny plots between the green and red maternal trees. Both trees are from the southern Sierra Nevada region.

**Fig. S5.**
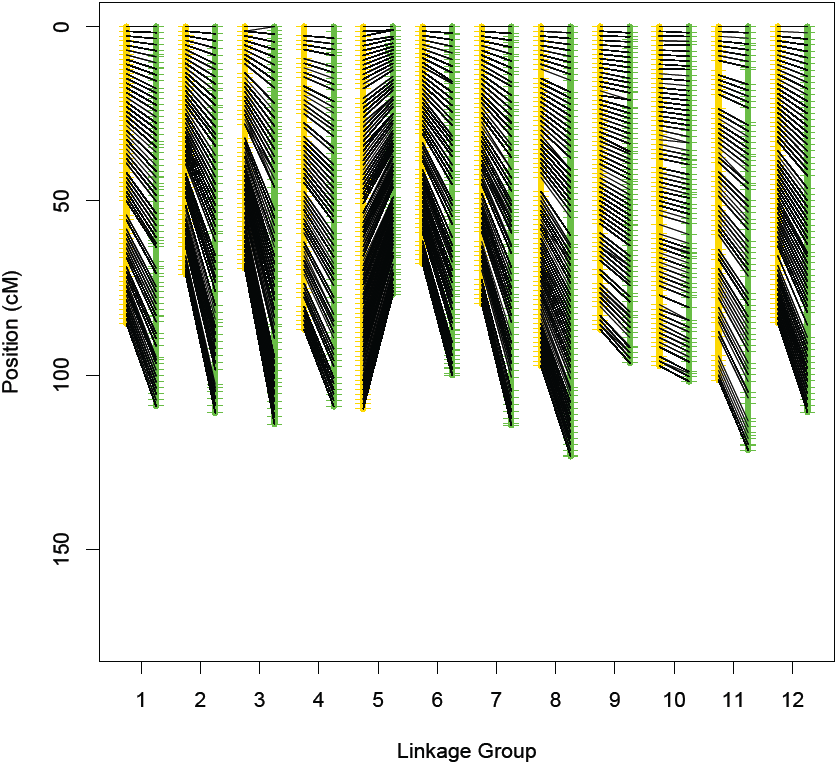
Pairwise synteny plots between the yellow (Klamath) and green (southern Sierra Nevada) maternal trees.

**Fig. S6.**
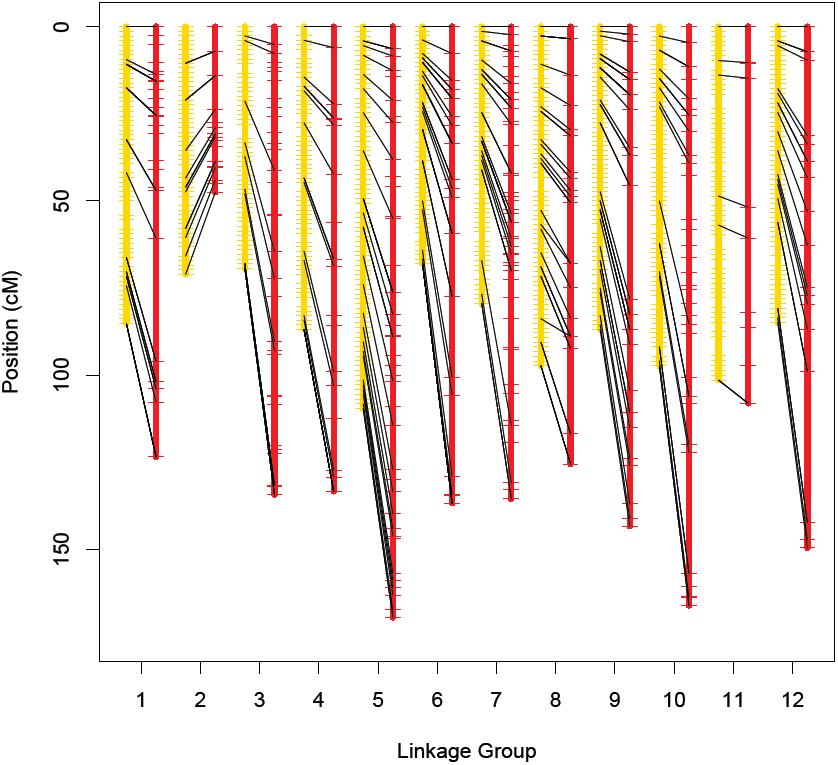
Pairwise synteny plots between the yellow (Klamath) and red (southern Sierra Nevada) maternal trees.

**Fig. S7.**
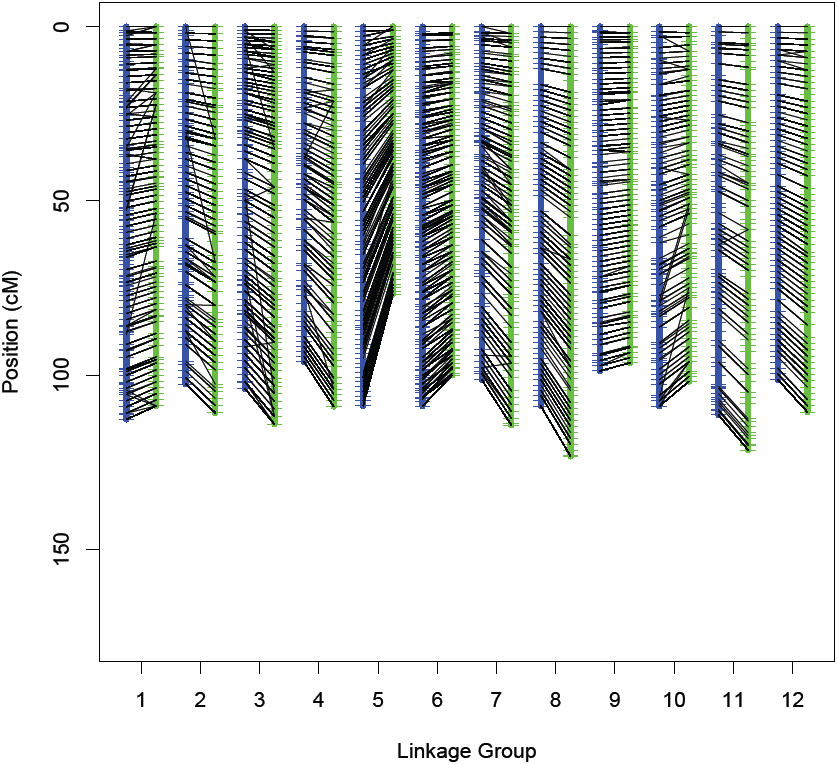
Pairwise synteny plots between the blue (Klamath) and green (southern Sierra Nevada) maternal trees.

**Fig. S8.**
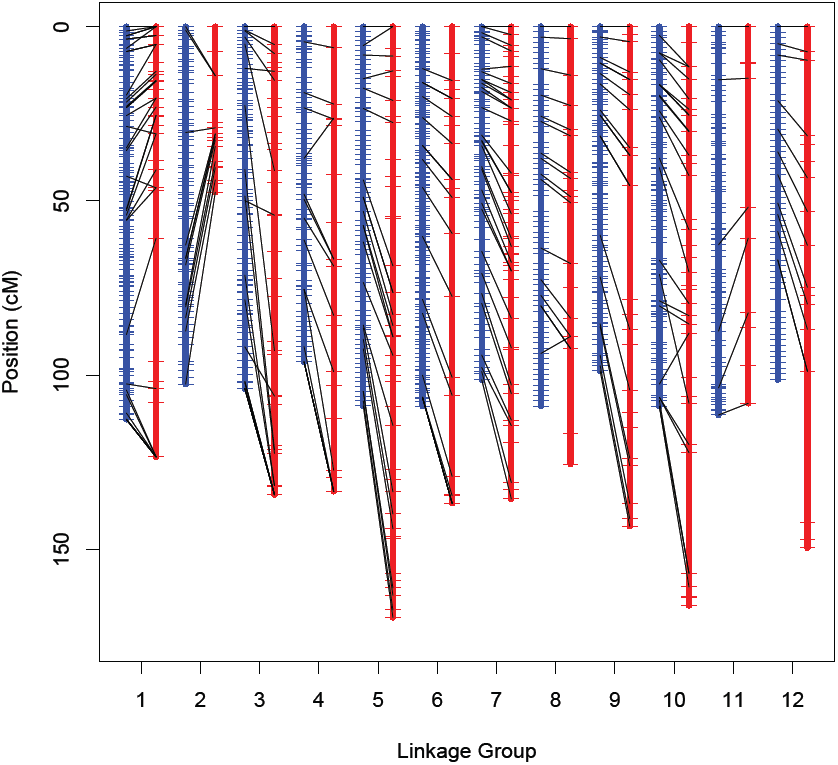
Pairwise synteny plots between the blue (Klamath) and red (southern Sierra Nevada) maternal trees.

**Fig. S9.**
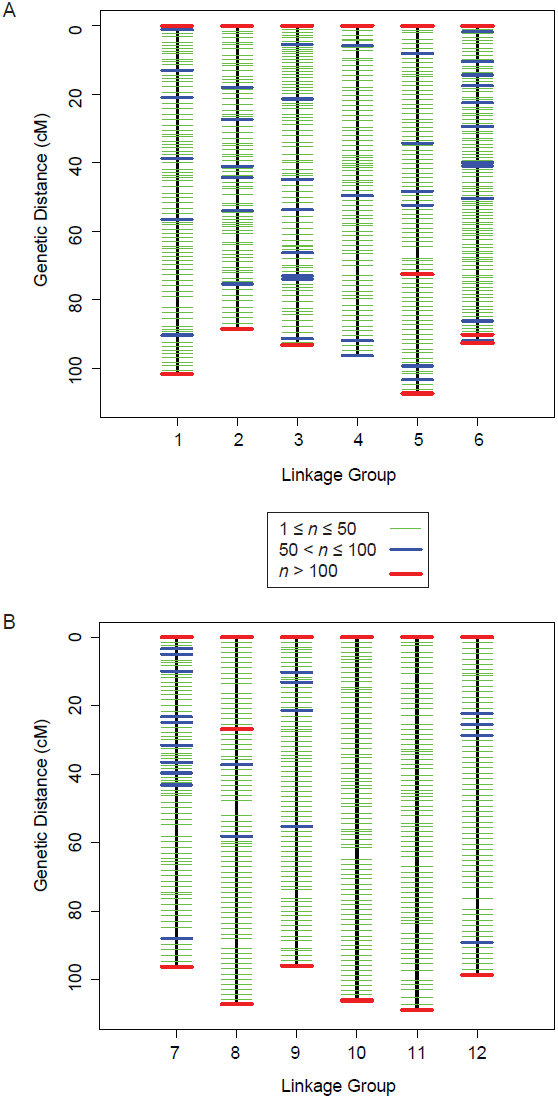
Consensus linkage map with positions colored by the number of contigs (n) mapped to each position. (A) Linkage groups 1 to 6. (B) Linkage groups 7 to 12. Additional information is given in Figure 3 within the main manuscript.

